# Nuclear α_1A_-Adrenergic Receptor Regulation of cAMP Production by an Inside-Out MAP Kinase Signaling Pathway in Cardiac Myocytes

**DOI:** 10.1101/2025.11.20.687712

**Authors:** Chase M. Fiore, Michael W. Rudokas, Riko Masuda, Brandon Khamsopha, Rinzhin T. Sherpa, Karni S. Moshal, Shailesh R. Agarwal, Robert D. Harvey

## Abstract

Sympathetic stimulation produces beneficial changes in cardiac function through β-adrenergic receptor (βAR) production of cAMP and subsequent alteration of electrical and mechanical activity. Long term activation of cAMP production also contributes to cardiac remodeling and detrimental changes associated with heart failure. However, sympathetic responses are mediated by the endogenous neurotransmitter norepinephrine (NE), which is also a potent α_1_-adrenergic receptor (α_1_AR) agonist, and α_1_AR activation can produce significant effects on the heart as well. What is less clear is how α_1_- and β-adrenergic responses interact with one another. Previous studies have demonstrated that α_1_AR activation can inhibit β-adrenergic regulation of electrical and mechanical activity of cardiac myocytes, although the signaling mechanisms involved were not previously known. In the present study, we used FRET-based biosensors in adult rat ventricular myocytes to demonstrate that this crosstalk effect involves inhibition of cAMP production by nuclear α_1A_ARs acting on βARs found on the plasma membrane. Furthermore, we established that this inside-out signaling mechanism involves a mitogen-activated protein kinase (MAPK) pathway that uncouples βARs from downstream signaling in a G protein coupled receptor kinase (GRK)/arrestin-dependent manner. These results reveal a novel, non-canonical signaling mechanism contributing to α_1_AR responses in the heart, and that this effect limits βAR production of cAMP by NE. This mechanism may contribute to the cardioprotective effect previously ascribed to α_1A_AR activation. These findings also clearly demonstrate the importance of considering the contributions of α_1_ and βARs together when studying the influence of the sympathetic nervous system on the heart.

## Introduction

Activation of the sympathetic nervous system results in the release of catecholamines that trigger a variety of cardiovascular responses [1]. In the heart, most of these effects are associated with the β-adrenergic receptor (βAR) signaling pathway, which involves stimulatory G protein (G_s_)-dependent activation of adenylyl cyclase (AC) and subsequent production of 3’,5’-cyclic adenosine monophosphate (cAMP) [2]. Increased levels of cAMP activate protein kinase A (PKA), which phosphorylates several key proteins involved in regulating the electrical and mechanical properties of cardiac myocytes [3]. However, cAMP production can also lead to the activation of the exchange protein activated by cAMP (Epac). Furthermore, chronic activation of Epac signaling pathways can contribute to pathological remodeling of the heart, cardiac arrhythmias, and heart failure [4].

While most cardiac responses to sympathetic stimulation are attributed to the activation of βARs, the primary neurotransmitter mediating these effects is norepinephrine (NE), which is also a potent α-adrenergic receptor (αAR) agonist. Furthermore, activation of α_1_ARs can elicit a variety of responses [5]. This includes effects believed to be cardioprotective [6,7]. These protective effects have been attributed, at least in part, to a mechanism involving α_1A_ARs found in the nucleus that are linked to a mitogen activated protein kinase (MAPK) signaling pathway that leads to activation of extracellular signal-regulated kinase (ERK) at the plasma membrane [8,9]. Yet, how this inside-out signaling mechanism produces its cardioprotective effects is less clear.

Activation of α_1_ARs can also modify cardiac myocyte responses to βAR stimulation [10–16], an effect that is often overlooked. Although the signaling mechanism responsible for this receptor crosstalk has not been identified, the fact that α_1_AR stimulation actually antagonizes βAR responses suggests that inhibition of cAMP production might be involved. If true, this mechanism could contribute to the cardioprotective effects that have been reported. The goal of the present study was to verify that α_1_AR regulation of βAR responses in adult ventricular myocytes does involve inhibition of cAMP production and determine whether the mechanism is mediated by α_1A_ARs acting through an inside-out MAPK-dependent signaling pathway.

## Materials and Methods

### Cell Isolation and Culture

Ventricular cardiac myocytes were isolated from 250-300 g adult Sprague-Dawley rats (Charles River, MA) of either sex, as previously described [17]. All protocols were performed in accordance with the Guide for the Care and Use of Laboratory Animals as adopted by the U.S. National Institutes of Health and approved by the Institutional Animal Care and Use Committee at the University of Nevada, Reno (protocol #20-05-1000, approval date 06/19/2023). In brief, rats were anesthetized by intraperitoneal injection of pentobarbital (150 mg/kg). The hearts were then excised, attached to a Langendorff apparatus, and perfused with a collagenase and protease containing solution to obtain isolated myocytes. These cells were then plated in M-199 media (Life Technologies, CA) supplemented with creatine (5 mM), taurine (5 mM), penicillin-streptomycin (1x), and bovine serum albumin (0.1%), and then transduced with an adenovirus encoding the Epac2-camps FRET biosensor. These cells were kept in culture (37°C and 5% CO_2_) for no more than 24 hours.

### Fluorescence Resonance Energy Transfer (FRET) Imaging

Live cell imaging was used to measure changes in cAMP activity detected by the Epac2-camps biosensor as previously described [17,18]. Briefly, cardiac myocytes expressing the biosensor were placed in a bath chamber on the stage of an Olympus IX71 inverted microscope and perfused with extracellular solution at room temperature. The donor fluorophore (eCFP) of the FRET based biosensor was excited using a Lambda DG-4 light source (Sutter Instruments, CA) with a D436/20 bandpass filter. Donor and acceptor (eYFP) fluorescence was measured simultaneously using an OrcaD2 dual chip CCD camera (Hamamatsu, Inc., Japan) fitted with 483/32 and 542/27 bandpass filters. Changes in whole cell cAMP activity were defined as the change in background and bleed-through corrected eCFP/eYFP fluorescence intensity ratio (ΔR) divided by the baseline ratio (R_0_). The FRET responses were then normalized to the maximum FRET response (MAX) generated by the application of 1 μM of the non-specific βAR agonist isoproterenol (ISO) and 100 μM of the non-specific phosphodiesterase inhibitor 3-isobutyl-1-methylxanthine (IBMX).

### Chemicals and Materials

G-protein receptor kinase inhibitor (CMPD101), pan PKC inhibitor (Gö 69683), forskolin (FSK), methoxamine (METH), corticosterone (CORT), prazosin (PRAZ), Raf inhibitor (SB 590885), MEK inhibitor (U0126), and norepinephrine (NE) were purchased from Tocris Bioscience (Bristol, UK). Lavendustin A was purchased from Cayman Chemical (Ann Arbor, MI). M-199 and penicillin-streptomycin were purchased from Life Technologies (Carlsbad, CA). All other chemicals were acquired from Sigma-Aldrich (St. Louis, MO).

### Statistical Analysis

Data values are represented as mean ± SEM of *n* cells from the hearts of *N* animals (*n/N*) in the text and figures. Cardiac myocytes from different isolations were randomly allocated to various experiments to reach an appropriate number of cells (n = 10-20) based on power calculations. The cutoff for statistical significance was set at p < 0.05. These p-values were calculated by ANOVA or paired two tailed t-test with correction for multiple comparisons, as needed, using Prism (version 10, GraphPad, CA).

### Data Availability

The datasets generated during the current study are available from the corresponding author on reasonable request.

## Results

### α1AR inhibition of βAR cAMP production

In previous studies, we have demonstrated that α_1_AR activation can antagonize βAR stimulation of cAMP-dependent ion channel responses in cardiac myocytes [8–14]. To verify that these actions are due to inhibition of cAMP production, as opposed to direct effects on ion channel activity, we measured responses in isolated adult rat ventricular myocytes (ARVMs) expressing the cAMP biosensor, Epac2-camps [19]. Exposure to 10 nM ISO resulted in a cAMP response that was 44 ± 4.3% (n/N = 13/3) of the maximum response observed upon exposure to 1 μM ISO plus 100 μM IBMX. Subsequent addition of the α_1_AR agonist methoxamine (METH, 3 μM) inhibited the ISO response by 67 ± 3.1% (n/N = 13/3) (Figure 1A & D). Methoxamine is a selective α_1_AR agonist. Consistent with this fact, in cells pre-treated with the α_1_AR antagonist prazosin (1 μM) [16,20] the inhibitory effect of METH was reduced to 23 ± 2.7% (n/N = 11/3) (Figure 1B & D). To determine if this effect involved α_1A_ARs, we repeated this experiment in cells treated with the α_1A_AR antagonist silodosin (1 μM) [21]. Under these conditions, the inhibitory effect of METH was reduced to 2.7 ± 1.6% (n/N = 11/3) (Figure 1C & 1D). These results support the idea that α_1_AR activation inhibits βAR production of cAMP in ARVMs, and that this effect specifically involves the α_1A_AR subtype.

**Figure 1:**
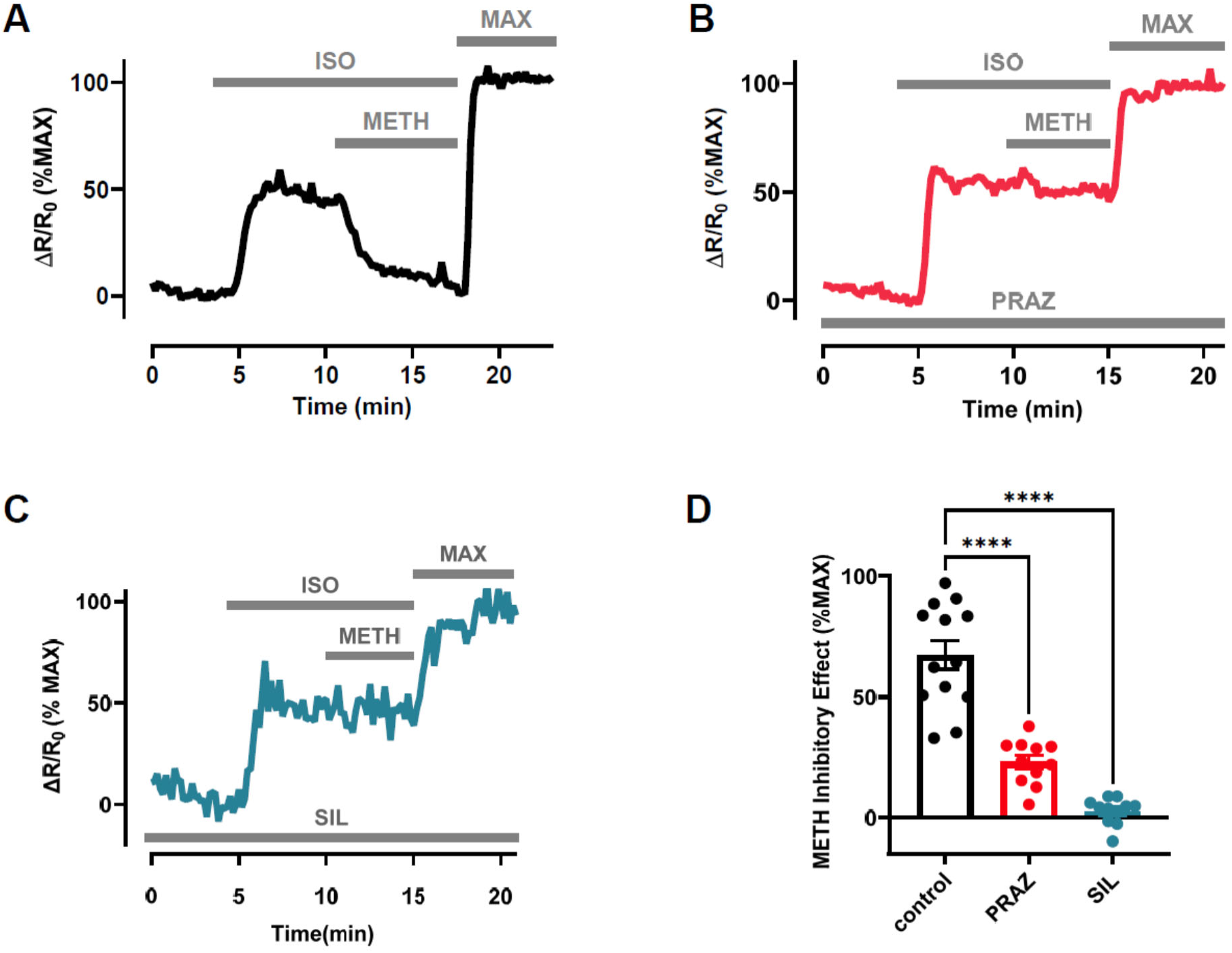
Effect of α_1_AR activation on βAR stimulated cAMP production. Time course of changes in cAMP activity detected by the Epac2-camps biosensor produced by exposure to the non-specific βAR agonist isoproterenol (ISO, 10 nM) followed by ISO plus the α_1_AR agonist methoxamine (METH, 3 μM) under control conditions **(A)** and in cells exposed to the α_1_AR antagonist prazosin (PRAZ, 1 μM) **(B)** or the α_1A_AR antagonist silodosin (SIL, 1 μM) **(C)**. FRET responses (ΔR/R_0_) were normalized to the magnitude of the maximum cAMP response (MAX) observed upon exposure to ISO (1 μM) + IBMX (100 μM) in the same cell. **(D)** Magnitude of METH-induced inhibition of the ISO response under control conditions (n/N = 13/3) was significantly reduced in the presence of PRAZ (n/N = 9/3, p < 0.0001) and SIL (n/N = 9/3, p < 0.0001), ANOVA with Dunnett’s multiple comparison test.

We then determined whether the ability of α_1_AR stimulation to inhibit cAMP production contributes to the net response produced by the endogenous sympathetic neurotransmitter norepinephrine (NE). While the effects of NE on the heart are often attributed to the activation of βARs alone, this neurotransmitter can activate both α and βARs. In untreated cells, exposure to 30 nM NE increased cAMP activity 15 ± 1.8% of maximum (n/N = 11/3) (Figure 2A & D). However, in the presence of 1 μM prazosin, the magnitude of the response to NE increased to 33 ± 3.3% of maximum (n/N = 11/3) (Figure 2B & D). Similarly, the cAMP response to NE was increased to 33 ± 3.1% of maximum (n/N = 10/3) (Figure 1C & D) in cells treated with silodosin. These results indicate that α_1A_AR attenuation of the cAMP response produced by βAR activation contributes to the net effect of the endogenous neurotransmitter NE.

**Figure 2:**
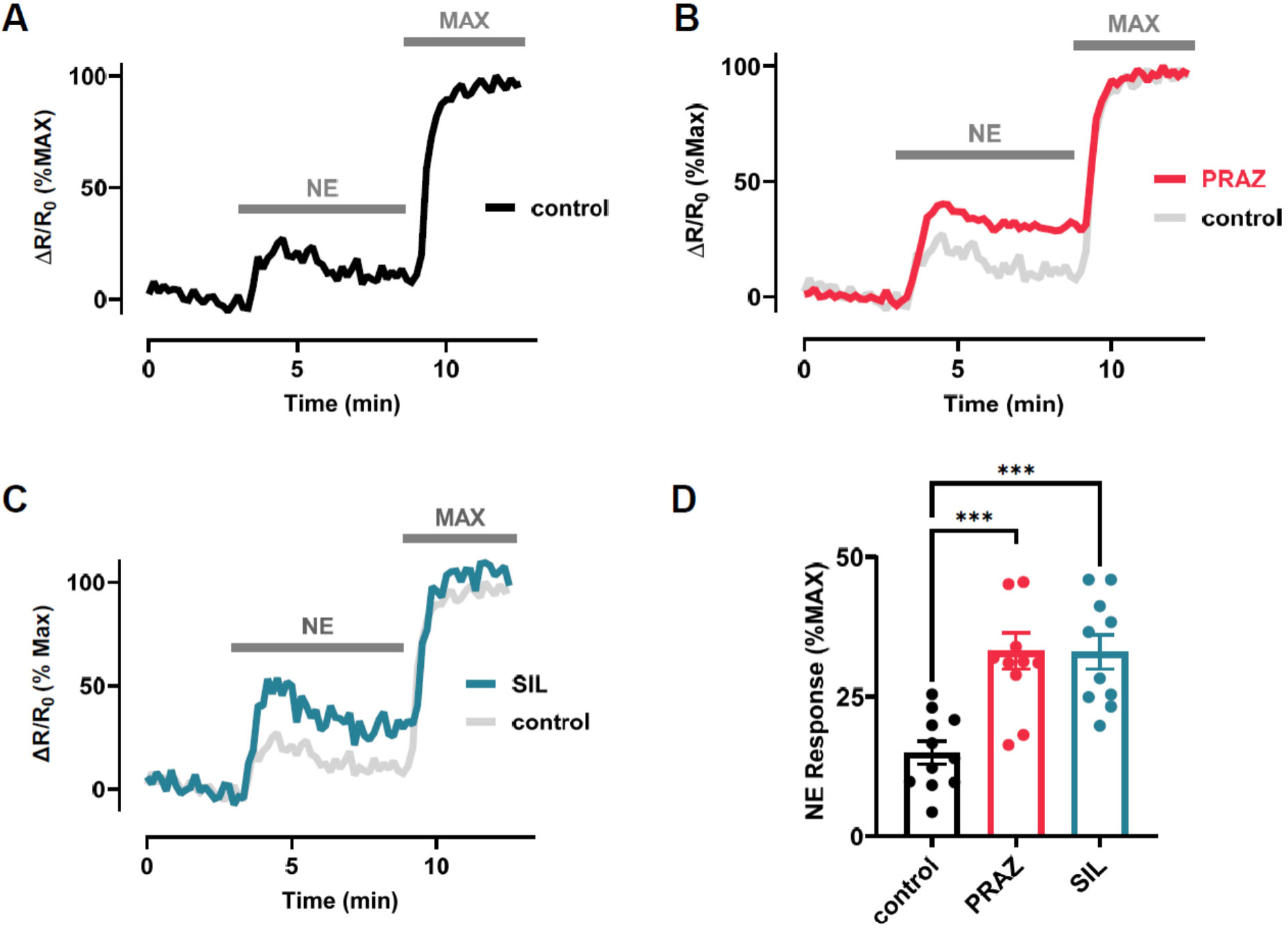
Contribution of α_1_AR stimulation to the cAMP response produced by the endogenous neurotransmitter norepinephrine (NE). **(A)** Time course of changes in cAMP activity detected by the Epac2-camps biosensor produced by exposure to NE (30 nM). **(B)** Time course of changes in cAMP activity produced by NE (30 nM) in the presence of the α_1_AR antagonist prazosin (PRAZ, 1 μM) and **(C)** the α_1A_AR antagonist silodosin (SIL, 1 μM). FRET responses (ΔR/R_0_) were normalized to the magnitude of the maximum cAMP response (MAX) observed upon subsequent exposure to ISO (1 μM) + IBMX (100 μM) in the same cell. **(D)** Average cAMP response produced by NE (n/N = 11/3, control) was significantly increased in the presence of PRAZ (n/N = 11/3, p = 0.0001) and SIL (n/N = 10/3, p = 0.0002), ANOVA with Dunnett’s multiple comparison test.

### Role of intracellular α_1_ARs

Previous studies using mouse ventricular myocytes have shown that the majority of α_1_ARs are located intracellularly, in and around the nucleus [8]. Using BODIPY-labeled prazosin, we determined that α_1_ARs are also found in and around the nucleus of rat ventricular myocytes (Figure 3C). To reach these receptors, agonists and antagonists must be able to cross the plasma membrane. In cardiac myocytes, endogenous agonists, such as NE, are known to enter the cell via the organic cation transporter (OCT3) [22]. To determine if α-adrenergic inhibition of βAR cAMP production involves activation of intracellular receptors, we conducted experiments in cells treated with corticosterone (CORT, 1 μM), which inhibits OCT3 [8,23]. Under these conditions, NE produced a cAMP response that was 44 ± 2.7% of maximum (n/N = 13/3) (Figure 3A & D). This is significantly larger than the NE response produced in untreated cells (see Figure 2). This is consistent with the idea that intracellular α_1A_ARs are responsible for inhibiting cAMP production. To confirm this conclusion, we repeated this experiment in cells treated with CORT (1 μM) and prazosin (1 μM). Previous studies have demonstrated that prazosin is able to diffuse across the plasma membrane, independent of OCT3 [8]. Under these conditions, the NE response was 44 ± 2.2 % of maximum (n/N = 14/3), which is not significantly different from the response observed in the absence of prazosin (Figure 3B & D), suggesting that CORT had prevented the ability of NE to activate intracellular α_1A_ARs. We also examined the response to METH in cells treated with 1 μM CORT. In these experiments, exposure to 10 nM ISO produced a cAMP response that was 49 ± 4.9% of maximum. Following addition of 3 μM METH, the cAMP response was 51 ± 3.1% (n/N = 4/2), which was not significantly different from the response to ISO alone (Supplemental Figure 1). These results support the conclusion that activation of intracellular α_1A_ARs is responsible for inhibiting βAR production of cAMP, and that both NE and METH rely on OCT3 to access these intracellular receptors.

**Figure 3:**
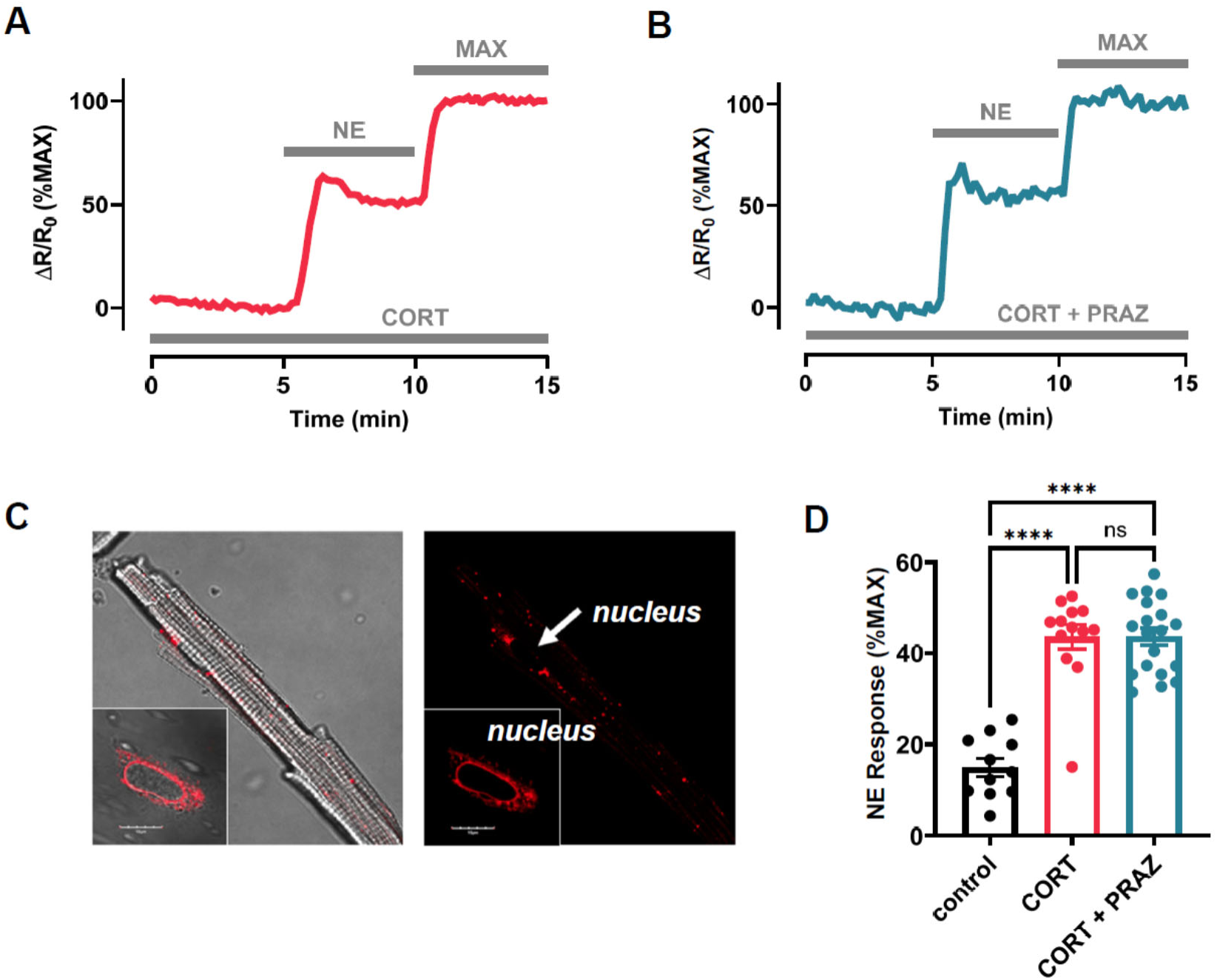
α_1A_AR inhibition of cAMP production depends on activation of nuclear receptors. **(A)** Time course of changes in cAMP activity detected by the Epac2-camps biosensor produced by exposure to NE (30 nM) in myocytes treated with the OCT3 inhibitor, corticosterone (CORT, 1 μM). **(B)** Time course of cAMP response to NE (30 nM) in myocytes treated with CORT (1 μM) plus PRAZ (1 μM). FRET responses (ΔR/R_0_) were normalized to the magnitude of the maximum cAMP response (MAX) observed upon subsequent exposure to ISO (1 μM) + IBMX (100 μM) in the same cell. **(C)** Fluorescence image of myocyte treated with BODIPY labeled prazosin (right panel). Nucleus denoted by white arrow. Merged brightfield and fluorescence images (left panel). Insets show labeling of isolated nuclei. **(D)** Average cAMP response produced by NE (n/N = 11/3, control) was significantly increased in the presence of CORT (n/N = 13/4, p < 0.0001) and CORT + PRAZ (n/N = 19/4, p < 0.0001). However, the response observed in the presence of CORT alone was not significantly affected by the addition of PRAZ (p = 0.99). ANOVA with Tukey’s multiple comparison test.

### PKC-dependent activation of Raf

The cardioprotective effect of α_1A_AR stimulation has been shown to involve a G_q_ signaling pathway [24], and nuclear α_1A_ARs have been shown to activate protein kinase C [25]. To determine if α_1A_AR inhibition of cAMP production is PKC-dependent, we examined the response to METH in cells treated with the PKC inhibitor Gö 6983 (5 μM) [26]. Under these conditions, exposure to 10 nM Iso produced a cAMP response that was 46 ± 4.4 % of maximum. However, subsequent addition of 3 μM METH did not significantly affect the cAMP response, which was 50 ± 4.6% of maximum (n/N = 14/3) (Figure 4A & B). This suggests that the α_1A_AR activation of PKC is involved in inhibition of cAMP production.

**Figure 4:**
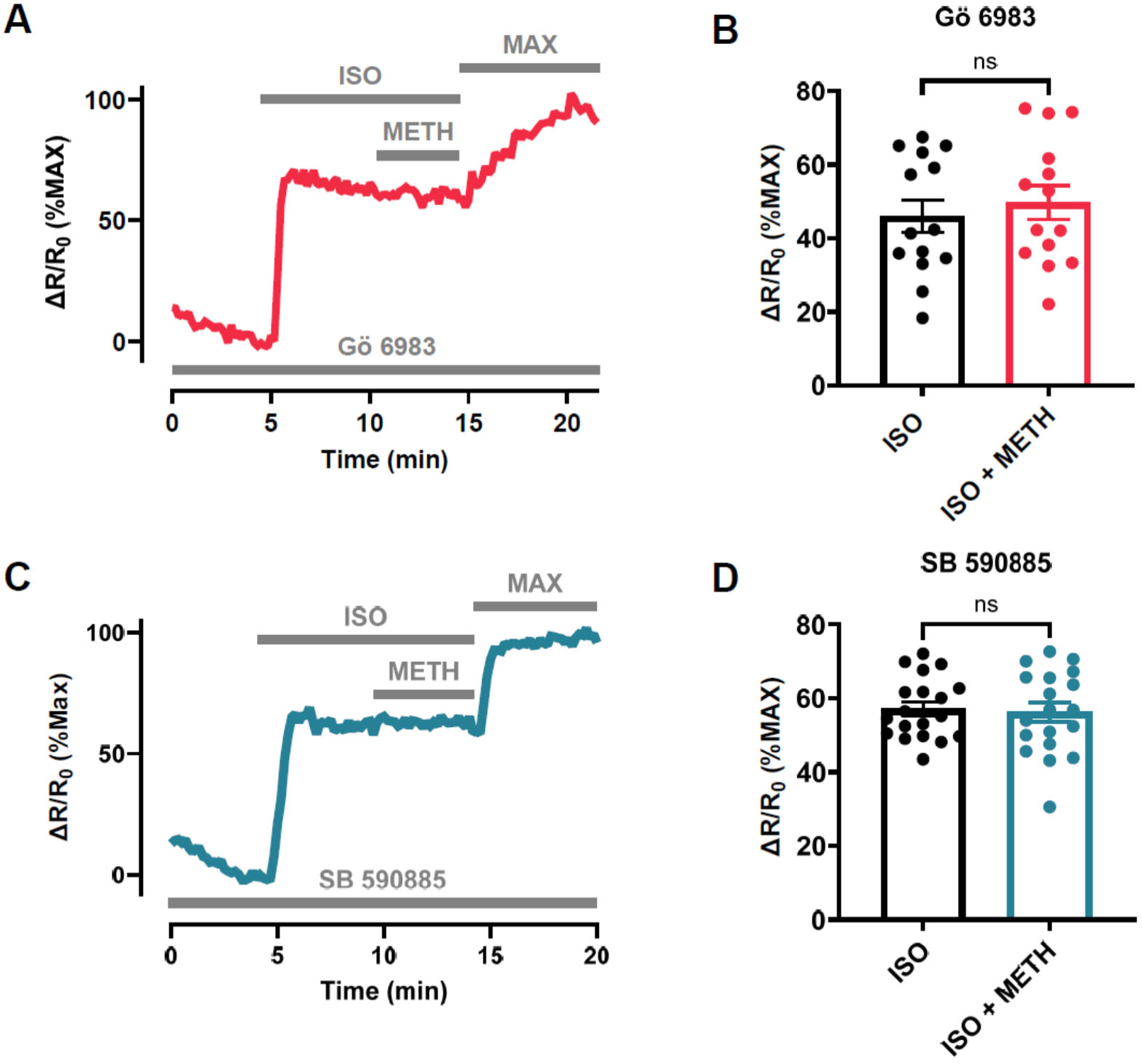
α_1A_AR inhibition of cAMP production involves activation of protein kinase C (PKC) and the serine-threonine kinase Raf. Time course of changes in cAMP activity detected by the Epac2-camps biosensor produced by exposure to isoproterenol (ISO, 10 nM) followed by subsequent addition methoxamine (METH, 3 μM) in the presence of the PKC inhibitor Gö 69683 **(A)** and or the Raf inhibitor SB 590885 **(C)**. FRET responses (ΔR/R_0_) were normalized to the magnitude of the maximum cAMP response observed upon exposure to ISO (1 μM) + IBMX (100 μM) in the same cell. The average response to ISO (10 nM) was not significantly affected by the addition of METH (3 μM) in cells treated with Gö 69683 (n/N = 14/4, p = 0.081) **(B)** or in cells treated with SB 590885 (n/N = 19/4, p 0.6653) **(D)** (paired t-tests).

We next tested the hypothesis that α_1A_AR regulation of cAMP production is linked to the MAPK signaling pathway through the activation of Raf. It has previously been shown that nuclear α_1A_ARs can activate ERK via MEK. Furthermore, MEK can be activated directly by PKC. However, MEK is more commonly activated by either Ras or Raf, and PKC is capable of directly activating Raf, independent of Ras [27]. To determine if Raf is part of the signaling pathway associated with the inhibition of cAMP production, we examined the response to METH in cells treated with the Raf inhibitor SB 590885 (10 μM) [28]. Under these conditions, 10 nM ISO produced a cAMP response that was 57 ± 1.9% of maximum (Figure 4C & D). However, following addition of 3 μM METH, the cAMP response remained at 56 ± 2.6% of maximum (n/N = 19/4). These results suggest that PKC activation of Raf is an integral part of the signaling pathway responsible for α_1A_AR inhibition of βAR cAMP production in cardiac myocytes (see Figure 7).

### MEK-dependent activation of ERK

We next looked specifically at whether MEK was involved in α_1A_AR inhibition of cAMP production by conducting experiments in cells treated with the MEK inhibitor U0126 (10 μM) [29]. Under these conditions, 10 nM ISO produced a cAMP response that was of 69 ± 3.7% of maximum. Following addition of 3 μM METH, the response remained at 69 ± 5.9%) (n/N = 11/4) (Figure 5A & B). These results further support the idea that α_1_AR inhibition of βAR production of cAMP involves a MAPK signaling pathway [8,9].

**Figure 5:**
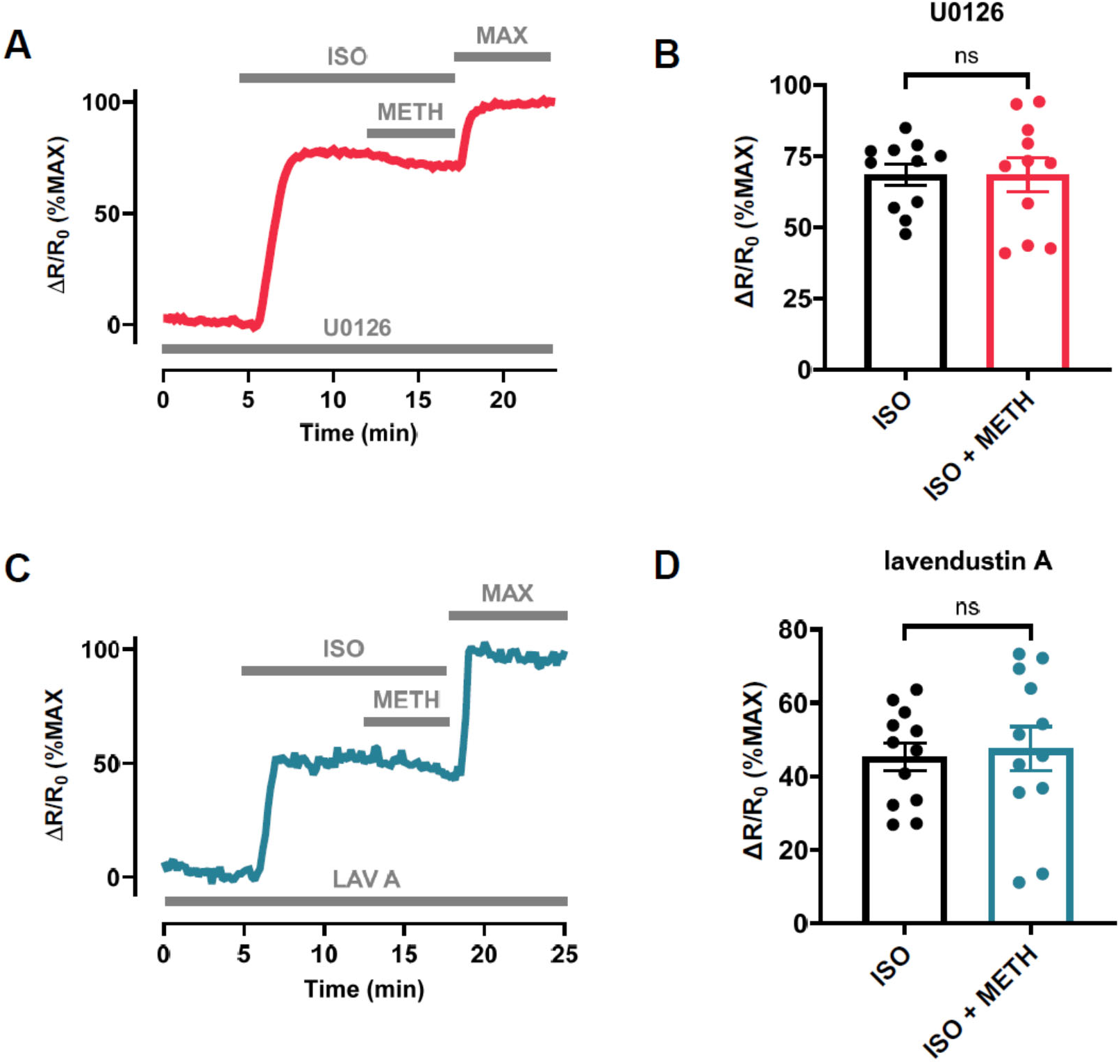
α_1A_AR inhibition of cAMP production involves mitogen activated protein kinase kinase (MEK) and tyrosine kinase activity. Time course of changes in cAMP activity detected by the Epac2-camps biosensor produced by exposure to isoproterenol (ISO, 10 nM) followed by subsequent addition methoxamine (METH, 3 μM) in the presence of the MEK inhibitor U0126 (10 μM) **(A)** and the tyrosine kinase inhibitor lavendustin A (LAV A) **(C)**. FRET responses (ΔR/R_0_) were normalized to the magnitude of the maximum cAMP response (MAX) observed upon exposure to ISO (1 μM) + IBMX (100 μM) in the same cell. The average response to ISO (10 nM) was not significantly affected by the addition of METH (3 μM) in cells treated with U0126 (n/N = 11/4, p = 0.99) **(B)** or in cells treated with LAV A (n/N = 12/3, p = 0.70) **(D)** (paired t-tests).

MEK is a dual specificity kinase that activates ERK by phosphorylating both threonine and tyrosine residues. Furthermore, we have previously demonstrated that α_1_AR inhibition of cardiac ion channel responses to βAR stimulation can be blocked by inhibiting tyrosine kinase activity [16]. To confirm that this same mechanism is also responsible for inhibition of cAMP production, the response to METH was first examined in cells treated with the tyrosine kinase inhibitor lavendustin A (5 μM) [16]. Exposure to 10 nM ISO produced an increase in cAMP activity that was 45 ± 3.8% of maximum (n/N = 12/3), but upon subsequent addition of 3 μM METH the response was 48 ± 6.1% (n/N = 12/3) indicating that the inhibitory effect had been blocked (Figure 5C & 5D). To confirm this result, we conducted a second set of experiments using cells treated with the tyrosine kinase inhibitor genistein (50 μM) [16]. Under these conditions, exposure to 10 nM ISO produced an increase in cAMP activity that was 66 ± 2.4% of maximum (n/N = 12/3). However, subsequent exposure to 3 μM METH did not significantly affect the size of the response to ISO, which was still 60 ± 2.9% of maximum (n/N = 12/3) (Supplemental Figure 3). These results are consistent with the idea that α_1A_AR inhibition of cAMP production involves tyrosine kinase-dependent activation of ERK (see Figure 7).

### G-protein coupled receptor kinase (GRK) dependent mechanism

Mitogen kinase signaling is commonly associated with the activation of ERK, which then translocates to the nucleus, where it regulates gene expression [30]. However, the cardioprotective effect of α_1_ARs has been linked to activation of ERK at the plasma membrane through an inside-out mechanism [8]. Although it has been suggested that this then affects mitochondrial function [31], ERK acting at the plasma membrane has also been shown to phosphorylate β-arrestin, increasing its activity and uncoupling receptors from downstream signaling [32]. Because β-arrestin can lead to uncoupling of βARs in the heart [33], we examined the possibility that this may be linked to α_1_AR inhibition or uncoupling of βAR production of cAMP. To test this hypothesis, we examined the response to METH in cells treated with the G-protein coupled receptor kinase (GRK) inhibitor CMPD101 (3 μM) [34,35]. Because GRK phosphorylation of βARs is required for β-arrestin binding, we hypothesized that inhibiting GRK would pre-empt any effect that ERK phosphorylation might have. Under these conditions, exposure to 10 nM ISO produced an increase in cAMP activity that was 52 ± 2.9% of maximum. Furthermore, following subsequent addition of 3 μM METH the response remained elevated at 54 ± 2.8% (n/N = 16/5) (Figure 6A & B), consistent with the idea that α_1_AR inhibition of βAR production of cAMP involves ERK phosphorylation of β-arrestin (see Figure 7).

**Figure 6:**
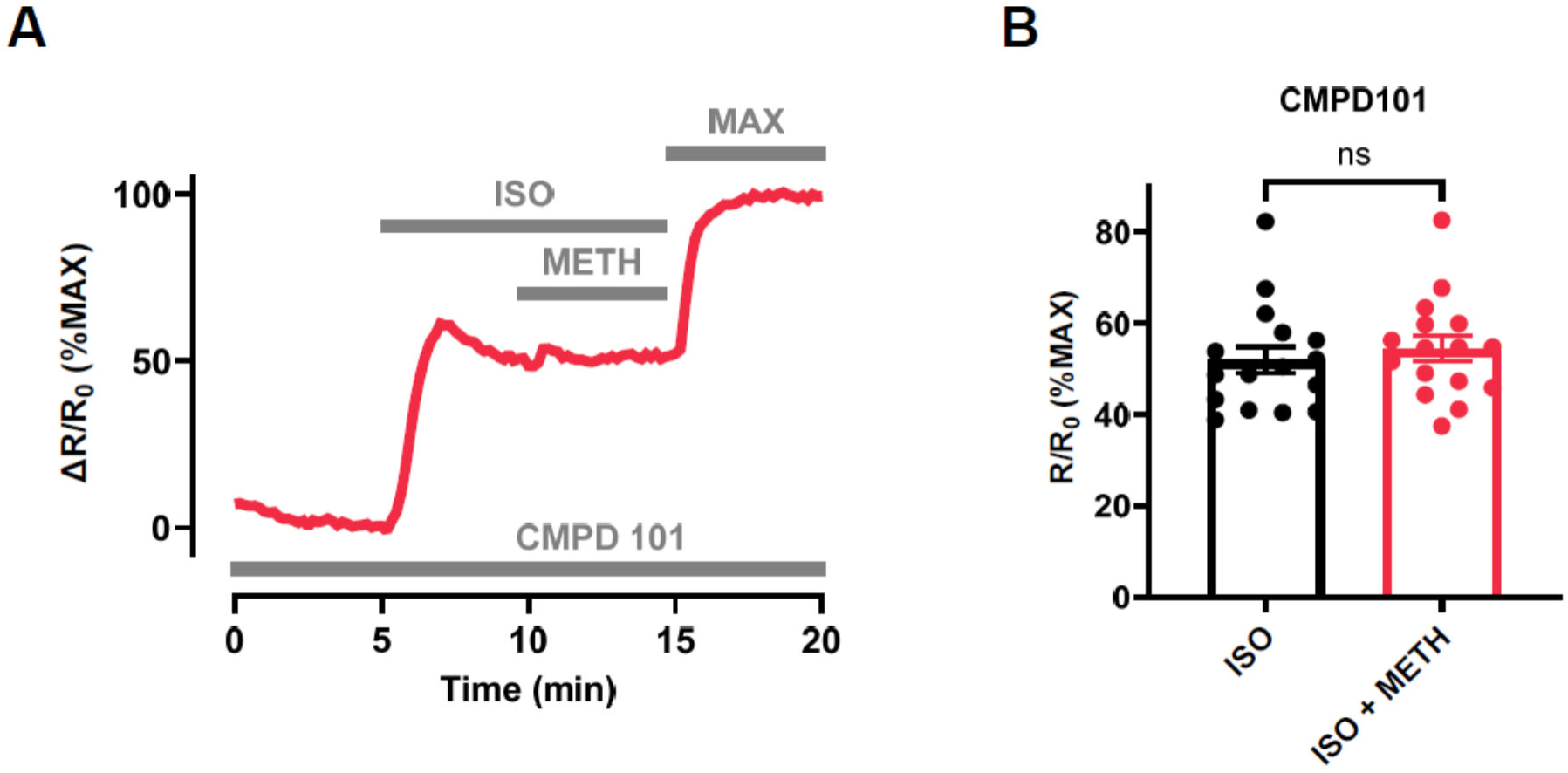
α_1A_AR inhibition of cAMP production depends on G protein coupled receptor kinase (GRK) activity. Time course of changes in cAMP activity detected by the Epac2-camps biosensor produced by exposure to isoproterenol (ISO, 10 nM) followed by subsequent addition methoxamine (METH, 3 μM) in the presence of the GRK inhibitor CMPD101 (3 μM) **(A)**. FRET responses (ΔR/R_0_) were normalized to the magnitude of the maximum cAMP response (MAX) observed upon exposure to ISO (1 μM) + IBMX (100 μM) in the same cell. The average response to ISO (10 nM) was not significantly affected by the addition of METH (3 μM) in cells treated with CMPD101 (n/N = 16/5, p = 0.13) (paired t-test).

**Figure 7:**
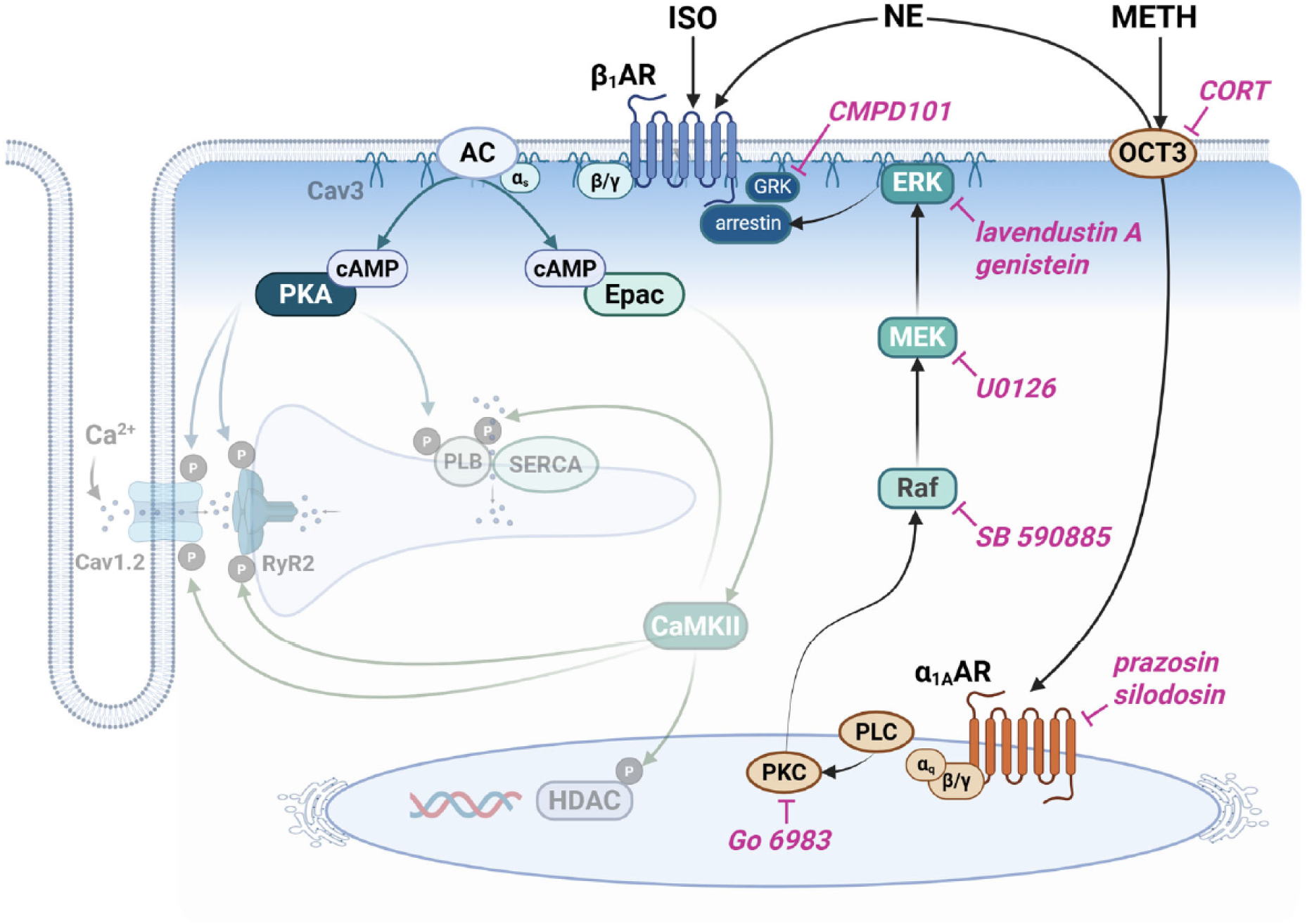
Inside-out signaling pathway responsible for α_1_AR inhibition of βAR cAMP production. α_1A_ARs in the nucleus inhibit cAMP production by βARs found at the plasma membrane by acting through a mitogen activated protein (MAP) kinase signaling pathway involving protein kinase C (PKC), Raf, and MEK. MEK activates ERK through a tyrosine kinase-dependent mechanism, and ERK can block βAR production of cAMP by activating arrestin, which depends on G protein kinase (GRK) phosphorylation of βARs. Activation of intracellular α_1A_ARs by norepinephrine (NE) and methoxamine (METH) requires uptake of these agonists by the organic cation transporter OCT3. Inhibiting each step in this pathway with the compounds indicated can antagonize the effect of α-agonists on βAR production of cAMP (see text for details). Created in BioRender. Harvey, R. (2025) https://BioRender.com/ksom4md

## Discussion

The sympathetic nervous system plays an essential role in regulating cardiac function through changes in heart rate and contractility. Chronic stimulation also contributes to changes in gene expression resulting in cardiac remodeling associated with heart failure [36,37]. These effects are mediated by the endogenous neurotransmitter NE, which activates both α- and βARs. Although the effects of NE on the heart are most commonly associated with β_1_AR activation, α_1_ARs can exert significant influence on cardiac function as well. This includes enhancing contractility, promoting adaptive hypertrophy, inducing ischemic preconditioning, and preventing myocyte death [5,6].

It is less clear how responses to α_1_- and β-adrenergic stimulation of the heart interact with one another. Previous studies have demonstrated that acute activation of α_1_ARs can inhibit βAR regulation of ventricular contraction as well as electrical activity [10–12,15,16,38]. One possible explanation for this behavior is that α_1_AR stimulation, acting through one of its canonical signaling pathways, exerts its influence at the level of the effector involved. Along these lines, it has been suggested that α_1_AR antagonism of cAMP-dependent L-type Ca^2+^ channel activity can be explained by a protein kinase C (PKC) dependent mechanism acting directly at the level of the ion channel [13]. However, this cannot explain the ability of α_1_AR stimulation to inhibit all β-adrenergic responses [15,16]. A simpler explanation would be that α_1_ARs somehow affect βAR production of cAMP. Several studies have examined the effect of α_1_AR activation on cAMP activity in various cardiac preparations using conventional biochemical assays, and the reported results have ranged from inhibition to stimulation [39–44]. The goal of the present study was to evaluate changes in cAMP activity using a newer, more sensitive approach. To that end, we measured changes in cAMP activity in adult ventricular myocytes expressing the FRET-based biosensor Epac2-camps, using a live cell imaging technique. The results of the present study clearly demonstrate that α_1_AR stimulation inhibits βAR production of cAMP. We also establish that this effect involves the α_1A_AR subtype. Furthermore, we show that the effect that α_1A_AR activation has on cAMP production contributes to the overall response to NE.

Consistent with our previous work, we found that the α_1A_AR responses are blocked by inhibiting tyrosine kinase activity [16]. We originally hypothesized that this effect might be explained by phosphorylation of tyrosine residues found on the βAR. Such an effect can uncouple β_2_ARs from downstream signaling in heterologous expression systems [45]. However, the results of the present study indicate that tyrosine phosphorylation is more likely occurring at an obligatory step in the MAPK signaling pathway [46]. Consistent with this conclusion, we demonstrated that the effect of α_1A_AR stimulation on cAMP production was blocked by inhibiting PKC, which can activate Raf. We also demonstrated that the α_1A_AR response could be blocked by directly inhibiting Raf, which can activate MEK. In addition, we found that we could block the α_1A_AR response by directly inhibiting MEK itself, which activates ERK through a mechanism that requires phosphorylation of tyrosine residues (Figure 7) [47].

The question then is how does activation of ERK affect βAR production of cAMP? As the name implies, extracellular signal-related kinase (ERK) is typically activated by events at the plasma membrane, which then produce changes in gene transcription in the nucleus [30]. However, it has been reported that α_1_ARs found in the nucleus of cardiac myocytes activate ERK in caveolar fractions of the plasma membrane, in an inside-out manner [8,9,25]. The results of the present study are consistent with α_1A_AR inhibition of cAMP production involving this inside-out pathway. This is supported by the fact that the response to METH was blocked by inhibiting OCT3, which indicates that this drug crosses the plasma membrane via the cation transporter, where it then activates intracellular receptors. We also found that inhibiting OCT3 enhanced the cAMP response to NE. This not only supports the idea that the α_1_ARs responsible for inhibiting cAMP production are intracellular, it also demonstrates that this inhibitory effect of α_1_AR stimulation contributes to the net cAMP response produced by NE.

It is worth noting that there have also been reports that NE can activate βARs found inside cardiac myocytes [23]. However, if the cAMP produced by NE in our experiments were due to activation of intracellular βARs, inhibition of OCT3 with CORT should have prevented this response. Our results can only be explained if the bulk of the cAMP produced in response to NE involves activation of extracellular βARs, and that these extracellular receptors are the ones being regulated through the ERK-dependent pathway (Figure 7). Although we did not determine which subtypes of βAR were affected by α_1A_AR stimulation, because NE is more selective for β_1_-receptors, our results suggest that this subtype is involved. However, we cannot rule out the possibility that a similar effect is occurring at β_2_-receptors as well.

The present results suggest that α_1_AR stimulation plays an important role in modulating cAMP responses in cardiac myocytes. This includes functional responses to the endogenous neurotransmitter NE observed under normal conditions. It is also conceivable that this pathway works to limit excess cAMP production in response to chronic βAR stimulation by the sympathetic nervous system under pathological conditions. In the Antihypertensive and Lipid Lowering Treatment to Prevent Heart Attack (ALLHAT) study, the α_1_AR antagonist doxazosin was found to significantly increase the incidence of cardiovascular events and it doubled the risk of heart failure in high-risk patients with hypertension. Similar evidence was obtained in the Vasodilator-Heart Failure Trial (V-HeFT), where patients with chronic heart failure were given various drugs to reduce afterload. Only those patients given prazosin failed to show improvement. Subsequent studies have demonstrated that α_1A_ARs activate ERK through an inside-out mechanism [8,9,25]. There is evidence that this is linked to the regulation of mitochondrial function [31]. However, inhibition of cAMP production could be another cardioprotective mechanism, providing benefits similar to those obtained with the use of βAR antagonists [37]. Consistent with this idea, we found that the effect on cAMP production appears to be mediated specifically by α_1A_ARs, the receptor subtype found to be cardioprotective [6]. The present findings further support the rationale for using selective α_1A_AR agonists as a treatment for heart failure [7,48].

Our results also demonstrate that the common practice of using the selective βAR agonist ISO is not the best approach for mimicking the effects of sympathetic stimulation on the heart. The influence of α_1A_ARs could have particularly important implications for understanding how sympathetic responses may change under conditions such as heart failure, where the ratio of α_1_ and β_1_ARs changes [48].

## Acknowledgements

The authors thank Maria Paz Salidias for help isolating cardiac myocytes. This work was supported by the National Institutes of Health National Heart, Lung, and Blood Institute Grants R01 HL145778, P01 HL164311, R01 HL161122, and P20 GM130459.

## Author Contributions

Study design (CMF, MWR, RDH) data acquisition (CMF, MWR, RM, BK, RTS, KSM), data analysis (CMF, MWR, RM, BK, RTS, KSM), data interpretation (CMF, MWR, SRA, RDH), and manuscript preparation (CMF, MWR, RDH)

### Abbreviations

AC: adenylyl cyclase;
α_1_AR: α_1_-adrenergic receptor;
ARVM: adult rat ventricular myocyte;
βAR: β-adrenergic receptor;
CORT: corticosterone;
FRET: fluorescence resonance energy transfer;
GEN: genistein;
IBMX: 3-isobutyl-1-methylxanthine;
ISO: isoproterenol;
LAV A: lavendustin A;
MAPK: mitogen-activated protein kinase;
METH: methoxamine;
NE: norepinephrine;
PKA: protein kinase A;
PKC: protein kinase C;
PRAZ: prazosin;
OCT3: organic cation transporter 3;
SIL: silodosin

