## Supplemental Figures for "Nuclear α_1A_-Adrenergic Receptor Regulation of cAMP Production by an Inside-Out MAP Kinase Signaling Pathway in Cardiac Myocytes"

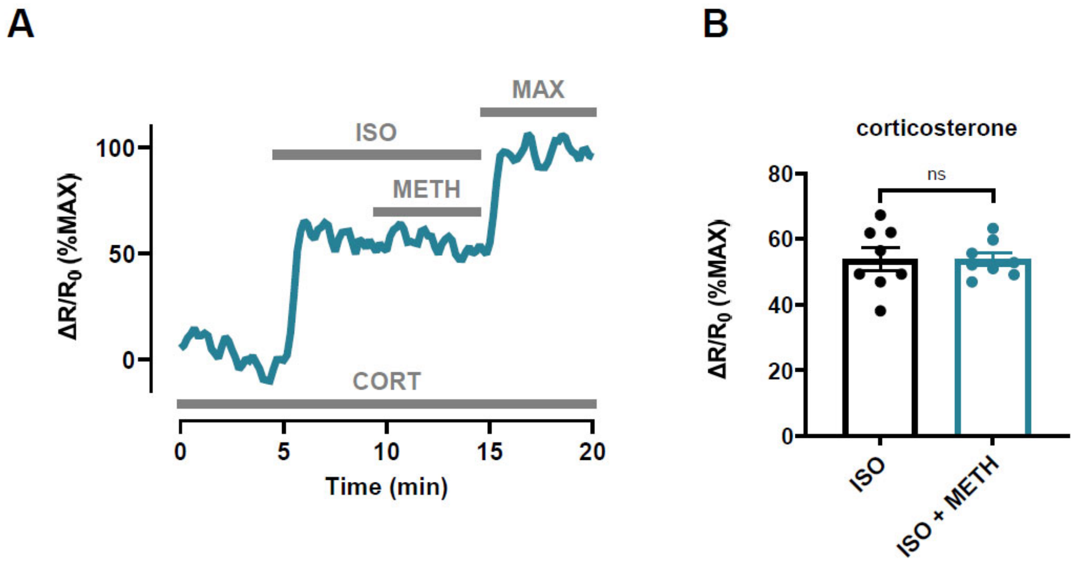


**Supplemental Figure 1:** $\alpha_{1A}$AR inhibition of cAMP production by the agonist methoxamine (METH) requires uptake by the organic cation transporter OCT3. **(A)** Time course of changes in cAMP activity detected by the Epac2-camps biosensor produced by exposure to isoproterenol (ISO, 10 nM) followed by subsequent addition methoxamine (METH, 3 μM) in myocytes treated with the OCT3 inhibitor, corticosterone (CORT, 1 μM). **(B)** The average response to ISO (10 nM) was not significantly affected by the addition of METH (3 μM) in cells treated with CORT (n/N = 8/3, $p = 0.98$) (paired t-test).

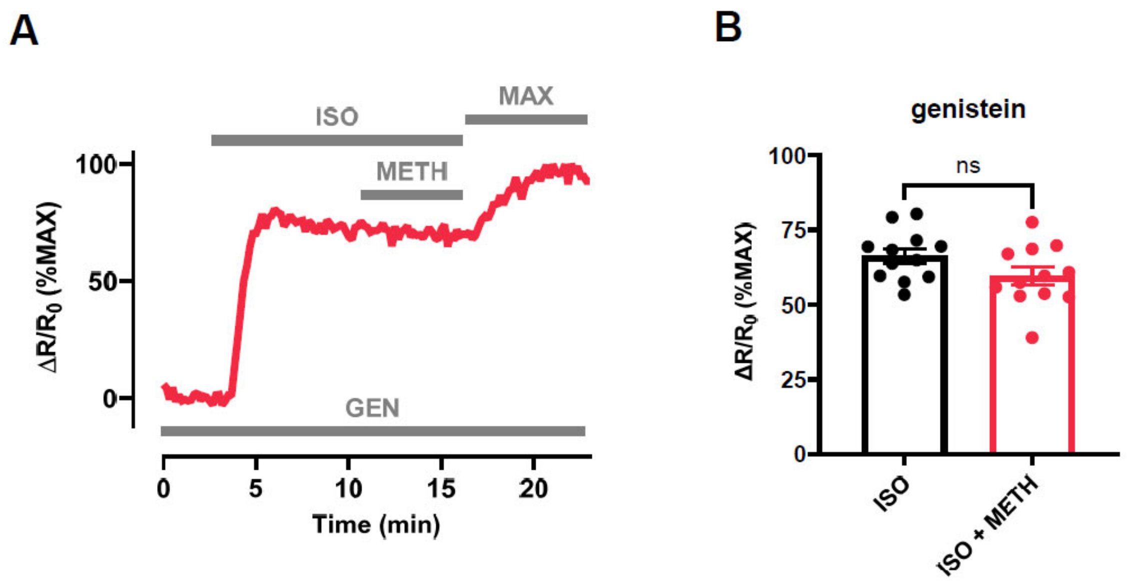


**Supplemental Figure 2:** $\alpha_{1A}$AR inhibition of cAMP production involves tyrosine kinase activity. **(A)** Time course of changes in cAMP activity detected by the Epac2-camps biosensor produced by exposure to isoproterenol (ISO, 10 nM) followed by subsequent addition methoxamine (METH, 3 μM) in myocytes treated with the tyrosine kinase inhibitor, genistein (GEN, 1 μM). **(B)** The average response to ISO (10 nM) was not significantly affected by the addition of METH (3 μM) in cells treated with GEN (n/N = 12/3, $p = 0.086$) (unpaired t-test).
